# The stress-protectant molecule trehalose mediates fluconazole tolerance in *Candida glabrata*

**DOI:** 10.1101/2024.07.16.603661

**Authors:** Qingjuan Zhu, Stefanie Wijnants, Regina Feil, Rudy Vergauwen, John E. Lunn, Mieke Van Ende, Patrick Van Dijck

**Author notes:** Laboratory of Molecular Cell Biology, Institute of Botany and Microbiology, KU Leuven, Leuven, Belgium.

## Abstract

The incidence of non-*albicans Candida* infections has witnessed a substantial rise in recent decades. *Candida glabrata (Nakaseomyces glabratus)*, an opportunistic human fungal pathogen, is accountable for both superficial mucosal and life-threatening bloodstream infections, particularly in immunocompromised individuals. Distinguished by its remarkable resilience to environmental stressors, *C. glabrata* exhibits intrinsic tolerance to azoles and a high propensity to swiftly develop azole resistance during treatment. The molecular mechanism for the high tolerance is not fully understood. In this work we investigated the possible role of trehalose in this tolerance. We generated mutants in the *C. glabrata TPS1*, *TPS2*, and *NTH1* genes, encoding trehalose 6-phosphate synthase (Tps1), trehalose 6-phosphate phosphatase (Tps2), and neutral trehalase (Nth1), respectively. As expected, the *tps1Δ* strain cannot grow on glucose. The *tps2*Δ strain demonstrated diminished trehalose accumulation and very high levels of trehalose 6-phosphate (T6P), the biosynthetic intermediate, in comparison to the WT strain. Whereas these higher T6P levels did not affect growth, the lower trehalose levels clearly resulted in lower environmental stress tolerance and a lower susceptibility to fluconazole. More interestingly, the *tps2Δ* strain completely lost tolerance to fluconazole, characterized by the absence of slow growth at supra-MIC concentrations of this drug. All these phenotypes are reversed in the *nth1*Δ strain, which accumulates high levels of trehalose. Our findings underscore the role of trehalose in enabling tolerance towards fluconazole in *C. glabrata*. We further show that the change in tolerance is a result of the effect that trehalose has on the sterol pattern in the cell, showing that accumulation of ‘toxic’ sterols correlate with absence of tolerance.

**Author summary:** *C. glabrata* is a yeast of significant medical importance, known for causing nosocomial outbreaks of invasive candidiasis. Its propensity to develop resistance to antifungal medications, notably azoles such as fluconazole, raises considerable concern. An underlying reason for the rapid development of resistance is its intrinsic tolerance to this drug. The underlying molecular mechanism of tolerance to fluconazole is heavily studied but not understood. This study sheds light on the involvement of trehalose in modulating tolerance to fluconazole. We have elucidated that trehalose serves not only as a protective agent against various stresses but also as a mediator of fluconazole resistance and tolerance. To start elucidating how this may work, we provide data that trehalose (or the enzymes affecting the amount of trehalose in the cell) alters the ergosterol type and level in the cells, thereby affecting tolerance.

## Introduction

Fungi infect billions of people every year, but still remain largely under-appreciated as human pathogens [1, 2]. Among candidemia cases, only five species, namely *C. albicans, C. glabrata, C. tropicalis, C. parapsilosis, and C. krusei,* account for 92% of the total [3, 4]. However, their distribution varies in population-based studies conducted in different geographical areas. *C. albicans* is the most encountered species, but there are significant differences in the number of cases caused by *C. glabrata* and *C. parapsilosis* [5]. Studies from Northern Europe and the USA have reported a high prevalence of candidemia caused by *C. glabrata* [3]. Isolates of this species are a threat to human health due to their ability to rapidly develop resistance to antifungal agents [6]. Unlike other *Candida* species that are diploid and usually require alterations in both alleles to confer resistance, *C. glabrata* is a haploid organism and only a single amino acid alteration may be required to confer resistance [7]. Therefore, greater effort is necessary to improve the available antifungals, as well as to find more potent and safer compounds with fungicidal action, or that prevent virulence attributes to be expressed. *C. glabrata* infections remain a clinical challenge, and there is an urgent need for new antifungal drugs with a novel mode of action to address the rising incidence of such infections [8].

Trehalose, characterized by its chemical structure as α-D-glucopyranosyl-(1,1)-α-D-glucopyranoside, is a non-reducing disaccharide consisting of two glucose moieties. Its significance in fungal conidia survival and germination is well-documented, primarily attributed to its role as a carbon source [9]. However, trehalose is also very important to provide protection against various stresses, including antifungal drugs [10, 11]. Trehalose is produced through a two-step enzymatic process involving trehalose-6-phosphate synthase (Tps1) and trehalose 6-phosphate phosphatase (Tps2). Initially, Tps1 catalyzes the conversion of uridine diphosphate (UDP)-glucose and glucose 6-phosphate into trehalose6-phosphate (T6P) and UDP. Subsequently, Tps2 converts T6P into trehalose and phosphate [12, 13]. Conversely, the breakdown of trehalose primarily occurs via hydrolysis facilitated by an α-glucosidase enzyme, which exhibits specificity towards trehalose as its exclusive substrate. This hydrolytic activity is carried out by trehalase enzymes. In *C. glabrata* these are encoded by *ATH1*, *NTH1*, and *NTH2* and we previously showed their importance in virulence in a gut colonization model [14]. The exploration of enzymes implicated in trehalose biosynthesis as potential antifungal targets is based on their absence in humans [15–17]. The significant role of trehalose in virulence has been demonstrated in various pathogens, both non-fungal and fungal. In non-fungal pathogens, trehalose biosynthesis has been found to enhance the pathogenicity of *Pseudomonas aeruginosa* in plants [18]. In fungal pathogens such as the opportunistic yeast *C. albicans* [19–21], the filamentous fungus *Aspergillus fumigatus* [22, 23], the phytopathogenic fungus *Magnaporthe oryzae* [24], and the meningitis-inducing basidiomycete *Cryptococcus neoformans* [25], trehalose also plays a crucial role in promoting virulence. These studies underscore the potential of trehalose metabolism as a promising avenue for antifungal intervention, offering insights into the vulnerabilities of diverse fungal pathogens [26]. Whereas *C. glabrata* is phylogenetically very close to *S. cerevisiae*, the role of trehalose metabolism in virulence of this pathogen has not yet been investigated. *C. glabrata* expresses one *TPS1* and one *TPS2* gene and, as mentioned above, three trehalase encoding genes [14].

In this work, we generated deletion strains in the trehalose biosynthesis genes *TPS1* and *TPS2*, and we also generated again an *NTH1* deletion strain, as the current background (*ATCC2001 his3Δ trp1Δ leu2Δ*) is different from that of the Van Ende study (14). We also generated strains where we expressed the *Cytophaga hutchinsonii TPSP* gene, a natural TPS–TPP bifunctional enzyme present in this bacterial species [27], into the wild type, *tps1*Δ and *tps2*Δ strains. This was undertaken to elucidate whether T6P or trehalose serves as the primary determinant for stress resistance as our hypothesis was that this enzyme does not produce any T6P but only trehalose. Our comprehensive analysis has revealed that the diminished intracellular trehalose levels, more than elevated T6P levels observed in the *tps2*Δ strain, correlate with increased susceptibility to thermal, oxidative, and osmotic stresses and a complete absence of tolerance to fluconazole. Conversely, the *nth1*Δ strain, which accumulates higher trehalose levels showed opposite phenotypes with an increase in tolerance towards fluconazole. Of particular interest is the observation that the difference in tolerance between the *tps2Δ* and *nth1Δ* strains may be caused by changes in the sterol types present in these two mutant strains, apart from a direct role that trehalose may play in stabilizing the membrane.

## Results

### The *C. glabrata TPS1* gene is required for growth on glucose as a carbon source

The Tps1 enzyme of *Saccharomyces cerevisiae* is required for growth on glucose as deletion of this gene results in uncontrolled influx of glucose into glycolysis, resulting in accumulation of sugar phosphates and concomitant depletion of ATP leading to apoptosis [28, 29]. In *C. albicans*, deletion of *TPS1* results in a growth defect on glucose, but only at temperatures above 39 °C [30]. To determine whether the Tps1 enzyme of *C. glabrata* is also involved in growth on glucose, we tested the different strains on SC medium supplemented with physiological glucose levels (5 mM) as well as with high glucose levels (100 mM). Our results show that the *TPS1* gene is essential for utilizing glucose as a carbon source, since the *tps1Δ* strain cannot growth in either liquid culture or on solid medium with glucose as the carbon source (Fig 1. A, B). Similar results (<u>Fig S1</u>) were obtained in the presence of trehalose or fructose as the carbon source. *C. glabrata* has a problem in using galactose as a carbon source and this was also clear in our experiments, where all strains showed a slow growth phenotype on galactose-containing media (<u>Fig S1</u>). Deletion of the *TPS2* or *NTH1* genes had no obvious effects on growth on any of the carbon sources in liquid or on solid media (<u>Fig S1</u> and <u>Fig S2</u>). To determine the role of the metabolites, we aimed to have strains with either low or high T6P or low or high trehalose. Based on studies in other species, we hypothesized that the *tps2Δ* strain accumulates high levels of T6P and low trehalose levels and that under optimal growth conditions, the wild type strain will have low T6P and low trehalose levels. In order to have a *C. glabrata* strain with no or low T6P but high levels of trehalose, we expressed the bifunctional T6P synthase/phosphatase enzyme (TPSP) of *Cytophaga hutchinsonii* [27] in the *tps1Δ* strain as the hyphothesis was that optimal flux of carbon in this enzyme results in a strain with low T6P and high trehalose levels. Expression of the *ChTPSP* gene in the *tps1*Δ strain restored growth on glucose, which was expected as the *ChTPSP* enzyme possesses both TPS and TPP enzymatic activities (Fig 1 A, B) [27].

**Fig 1.**
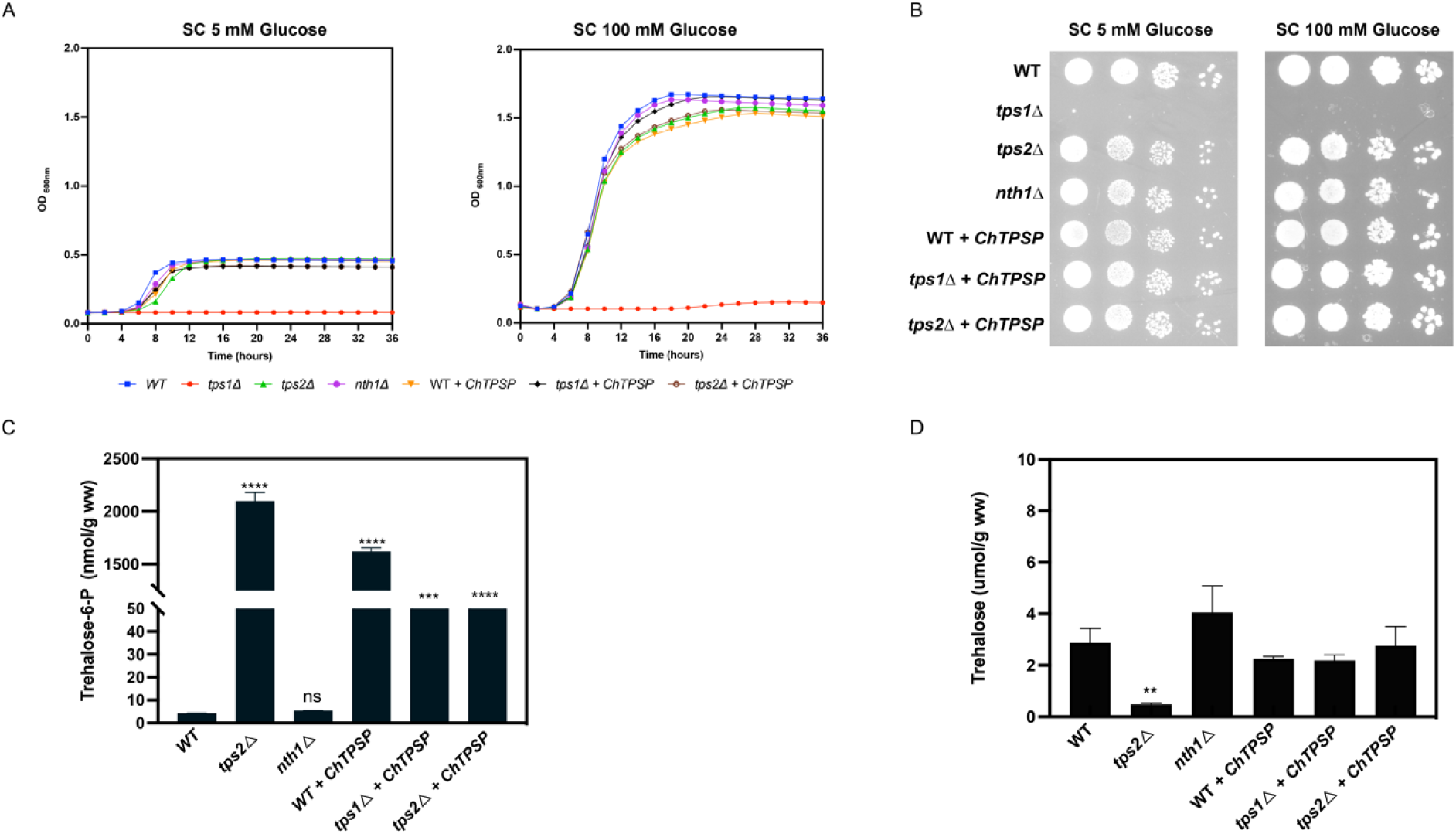
Deletion of *TPS1* affects the utilization of glucose as a carbon source. *C. glabrata* cells (WT, *tps1Δ*, *tps2Δ*, *nth1Δ, WT+ChTPSP, tps1Δ* +*ChTPSP* and *tps2Δ + ChTPSP* were grown at 37 °C in liquid medium and growth was monitored using a Multiskan microplate photometer. (A) The different strains were grown in SC medium supplemented with 5 mM glucose or 100 mM glucose. The OD_600nm_ was followed over time for 48 h. (B) The various strains were grown on solid SC medium supplemented with 5 mM glucose or 100mM glucose. The plates were incubated for 48 h at 37 °C before pictures were taken. (C) T6P levels of the different strains growing till exponential phase in SC plus 100 mM glucose at 37°C, 200 rpm; (D) trehalose levels of the different strains grown till exponential phase in SC plus 100 mM glucose at 37°C, 200 rpm. Average T6P and trehalose levels with SEM are shown of two experiments using three independent transformants of each strain. Statistical analysis was conducted by two-way ANOVA with Bonferroni correction; ns, no significant, *, P < 0.05; **, P < 0.01; ***, P < 0.001****, P < 0.0001.

### Expression of the bifunctional *C. hutchinsonii TPSP* gene still results in high levels of T6P upon expression in the *tps1Δ* strain

To understand the role played by Tps1 or Tps2, we needed to determine the levels of T6P and trehalose as these metabolites may contribute to the phenotype of the mutants. As the *tps1Δ* strain does not grow on glucose, we cannot use that strain directly. Therefore, we expressed a bifunctional enzyme with the aim to obtain a strain that has little T6P, as we expect efficient conversion of T6P into trehalose in this enzyme, but high levels of trehalose. However, expression of *ChTPSP* in the *tps1Δ* mutant resulted in only the same level of trehalose as observed for the wild type and a *tps2Δ* strain expressing the bifunctional enzyme (Fig 1D). Unexpectedly, expression of the bifunctional enzyme in the *tps1Δ* strain resulted in high levels of T6P (Fig 1E). These levels were still much lower than those obtained for the *tps2Δ* strain. As we previously observed in *C. albicans*, deletion of *TPS2* still results in some accumulation of trehalose. Deletion of *NTH1* resulted in increased trehalose levels as expected, and it had no effect on T6P levels. Based on these results, the remainder of the manuscript will focus on the wild type, the *tps2Δ* strain and the *nth1Δ* strain as these strains have either normal, high or low T6P and trehalose levels.

### Trehalose levels are important for heat, oxidative, and osmotic stress tolerance

Inside the human body, fungal pathogens are continuously exposed to different types of environmental stresses [31]. As trehalose is a disaccharide important for stress resistance [32], we assessed the growth phenotype of the deletion strains upon different stress treatments: oxidative stress (H_2_O_2_), salt stress (NaCl) and heat stress (39°C and 42°C). Heat stress of 42°C is too high for the human body but was chosen in accordance with previous studies, where this temperature was often used as test condition for heat stress [33]. The wild type can clearly not grow at this temperature but the *nth1Δ* strain still grows, albeit slowly, suggesting that the high amount of trehalose in these cells protects the cells against this stress (Fig 2). An opposite trend was observed under oxidative stress conditions, but the difference between the wild type and the *nth1Δ* strain was minor. We did not observe any differences between strains under osmotic stress conditions (Fig 2). The *tps2Δ* strain shows lower tolerance at 42 °C compared to the wild type strain and also a clearly lower tolerance towards oxidative and salt stress conditions (Fig 2).

**Fig 2.**
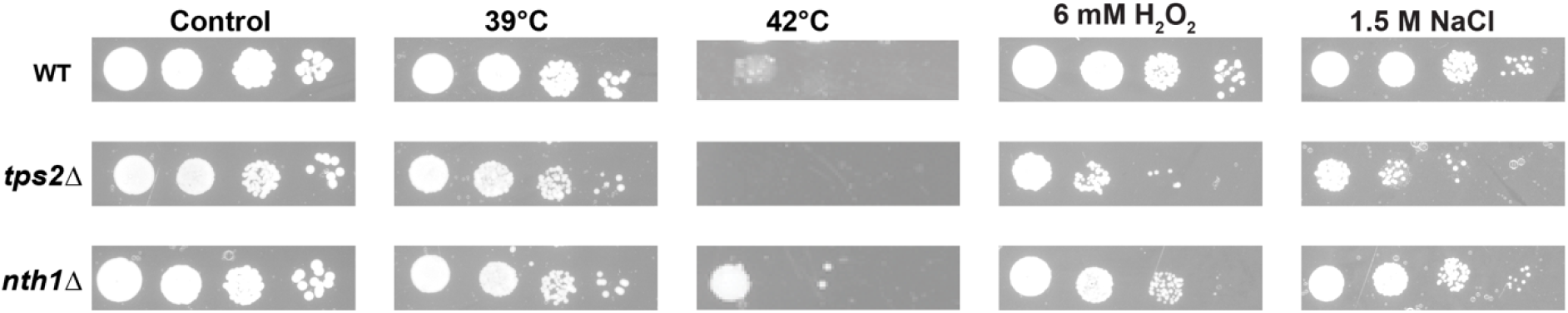
Trehalose provides tolerance against heat stress and is required for oxidative and salt stress tolerance. The different strains were grown overnight in SC medium with 100 mM glucose at 37°C. After three washing steps, 10-fold dilutions of cells were spotted on SC plus 100 mM glucose agar plates challenged with high temperature stress (at 39°C or 42°C), or with oxidative (H_2_O_2_) or salt (NaCl) stress and imaged after 48 hours of growth.

### Trehalose Accumulation Suppresses Resistance and Tolerance to Fluconazole in *C. glabrata*

Trehalose metabolism enzymes are interesting antifungal drug targets, and several groups are trying to find drugs that target these enzymes, but as far as we know, no compound targeting these enzymes is in any clinical trial [17, 34, 35]. Such inhibitors could also work synergistically with known antifungals as it was already shown in *C. albicans* that trehalose-deficient mutants are sensitive to amphotericin B and micafungin [21]. In *C. glabrata*, we did not observe a difference in sensitivity to amphotericin B between the WT and the *tps2Δ* or *nth1Δ* strains (Fig. S3). Unlike other typical *Candida* species, *C. glabrata* inherently exhibits reduced susceptibility to azole drugs, particularly fluconazole. To determine a possible role for trehalose metabolism in this fluconazole tolerance or resistance, we tested the growth of WT, *tps2*Δ, and *nth1*Δ strains in the presence and absence of fluconazole (Fig 3A). Deletion of *TPS2* resulted in strains that are very sensitive to fluconazole treatment, while the absence of the *NTH1* gene resulted in improved growth under fluconazole stress conditions.

**Fig 3.**
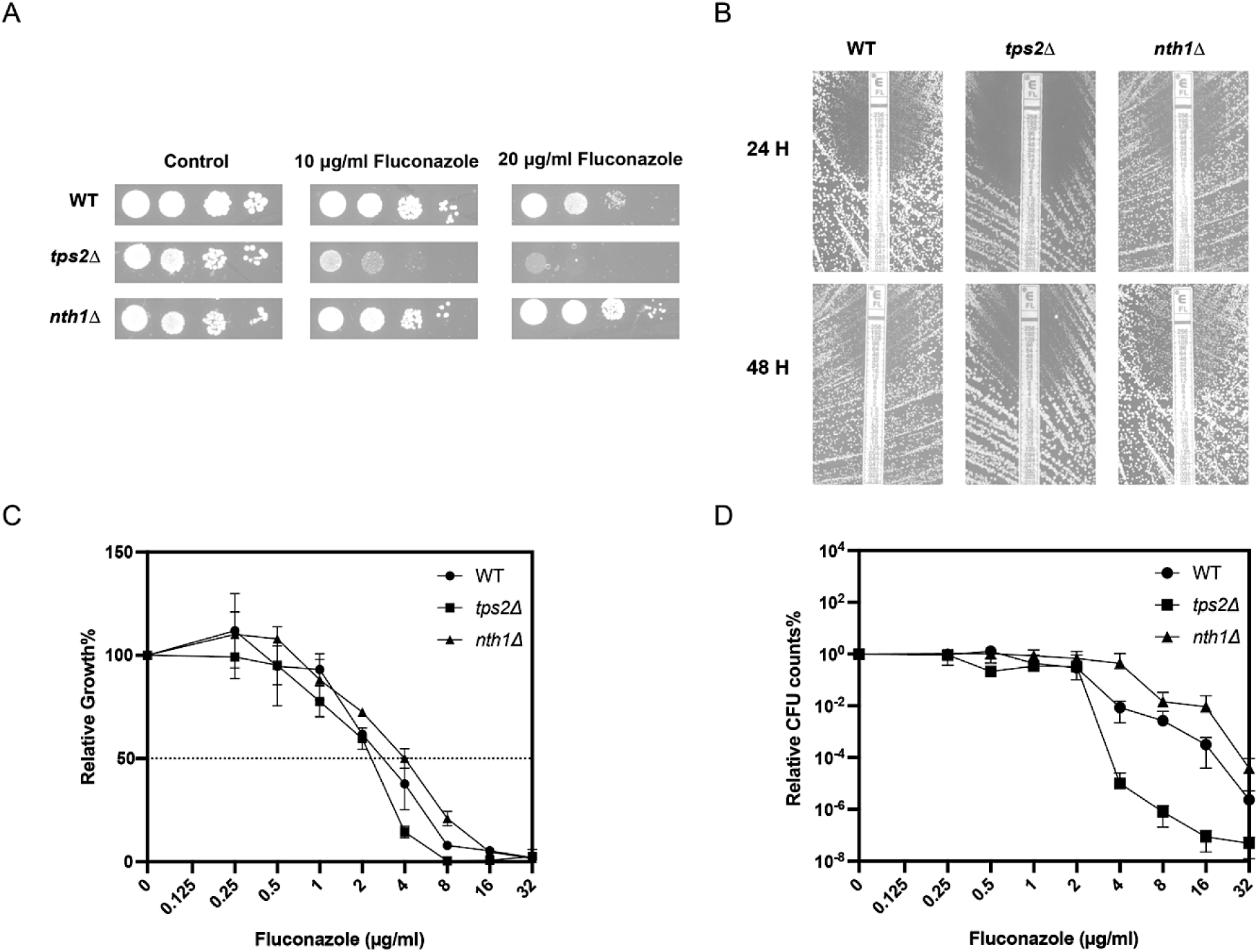
Trehalose plays an important role in tolerance against fluconazole. (A) Serial dilutions of the WT, *tps2Δ* and *nth1Δ* strains spotted on SC glucose plates. Pictures were taken after 48 h of incubation at 37°C. (B) Etest analysis at 37°C (24 and 48 hours) showing a “halo” in regions of the medium with high concentrations of fluconazole where the cells are unable to grow. (C) Growth profiles of mutant strains relative to the WT strain in a broth dilution assay (BDA) in the presence of different concentrations of fluconazole. (D) Tolerance assay. Data reflect the relative percentage of colony forming units from the WT and mutant strains after 48 hours of incubation at 37°C. The experiment was conducted two times with three biological and three technical repeats, and representative results are shown.

The minimum inhibitory concentration of fluconazole (MIC_flu_) for these strains was determined through Epsilometer test (Etest) analysis (Fig 3B) and broth microdilution assays (Fig 3C, D). From the Etest results, it can be inferred that the absence of the *NTH1* gene results in a decreased susceptibility to fluconazole, while deletion of the *TPS2* gene renders the cells more susceptible to fluconazole. The MIC values we obtain here are 8 µg/mL for the wild type, 2 µg/mL for the *tps2Δ* strain and 12 µg/mL for the *nth1Δ* strain. These data differ from the results obtained using the broth microdilution assay where the difference between the strains is less clear with MIC_50_ of 4 µg/mL for wild type, 2-4 µg/mL for *tps2Δ* and 4-8 µg/mL for the *nth1Δ* strain. Although the broth microdilution assay does provide an estimate of MIC_50_ value, it more importantly enables us to determine tolerance, based on 48 hours of growth on supra-MIC levels of fluconazole. Fig 3D shows that there is a strong difference in colony forming unit (CFU) counts at higher concentrations of fluconazole. The *tps2Δ* strain shows a strong reduction in tolerance, whereas the *nth1Δ* strain shows an increase in tolerance. This difference in tolerance is also clear from the Etest result in Fig 3B, where we see slow growth inside the halo of the *nth1Δ* strain and no growth inside the halo of the *tps2Δ* strain, after 48 hours of growth.

We additionally quantified the intracellular levels of trehalose and T6P to elucidate whether trehalose or T6P influences the resistance and tolerance to fluconazole in *C. glabrata*. In the analyzed strains, following a 60-minute exposure to fluconazole, both the WT and *nth1*Δ strains accumulated trehalose in response to the antifungal challenge, with an increased accumulation observed in the *nth1*Δ strain especially upon fluconazole treatment. However, there was no change in T6P levels observed in any of the three strains following fluconazole treatment.

### Impact of Trehalose Accumulation on Residual Ergosterol Levels and *ERG11* Gene Expression

Fluconazole targets Erg11, leading to an accumulation of lanosterol and a decrease in ergosterol levels in the presence of the drug [36]. This condition also prompts the conversion of lanosterol to 14-methylfecosterol and ultimately to the toxic compound 14α-methylergosta-8,24(28)-dienol, which constitutes a crucial aspect of the drug’s mode of action [37, 38]. To explore the potential involvement of ergosterol in mediating the impact of trehalose metabolism-related enzymes on fluconazole susceptibility, we assessed *ERG11* gene expression levels and ergosterol levels in the mutant strains. As illustrated in Fig 5, in the absence of fluconazole, there was only a slight difference observed between the control and mutant strains regarding lanosterol and ergosterol levels. Upon deletion of the *NTH1* gene, an increase in ergosterol levels was observed, whereas the *tps2*Δ strain exhibited decreased ergosterol levels. Furthermore, there was no production of 14α-methyl fecosterol and toxic sterol (14α-methylergosta-8,24(28)-dienol) observed. Upon fluconazole treatment, however, the proportion of remaining ergosterol in the *nth1*Δ strain was higher than in the control, while the *tps2*Δ strain displayed a significant decrease compared to the WT strain. Remarkably, we observed that absence of the *NTH1* gene resulted in less 14α-methyl fecosterol and toxic sterol formation compared to the control strain. Conversely, the absence of *TPS2* caused a lot of 14α-methyl fecosterol and toxic sterol formation. In summary, it appears that elevated trehalose levels correspond to increased ergosterol production, while reduced trehalose levels correlate with decreased ergosterol synthesis and higher levels of toxic sterols. These data can explain the difference in tolerance seen in Fig 3.

**Fig 4.**
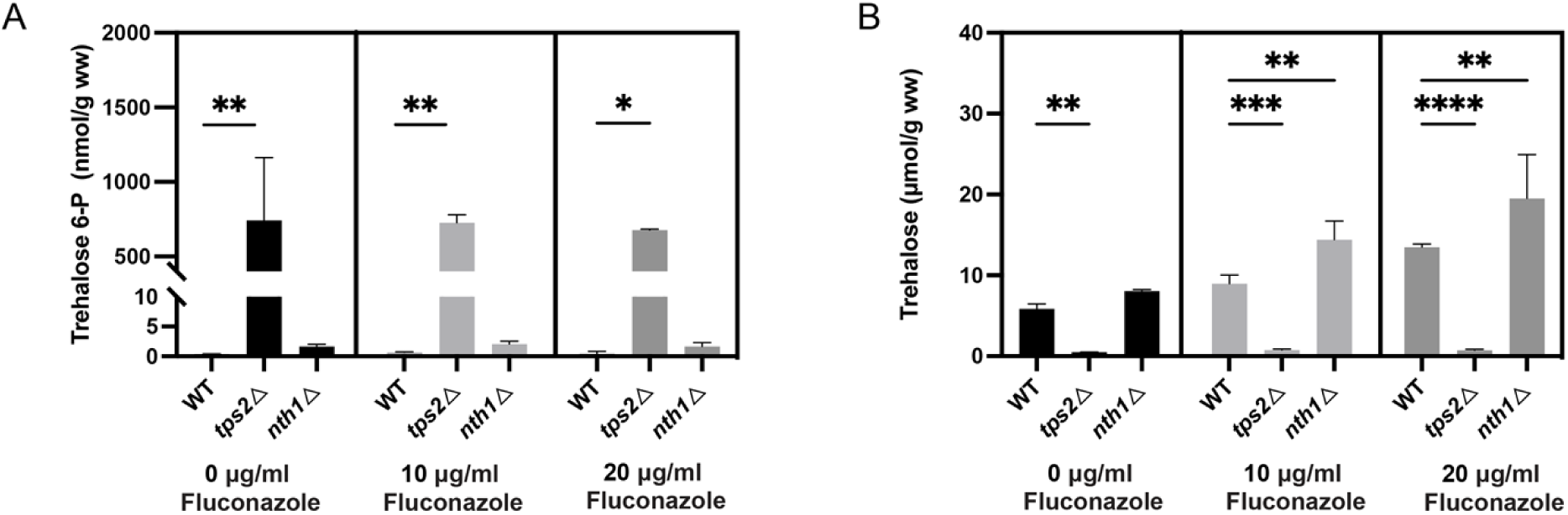
Intracellular T6P and trehalose levels in response to fluconazole stress. (A) T6P levels after fluconazole treatment. The cultures were divided in three and incubated further at 37°C with or without fluconazole for 60 minutes. (B) Trehalose levels in the presence and absence of fluconazole. The cultures were divided in three and incubated further at 37°C with or without fluconazole for 60 minutes (ww, wet weight). Average T6P and trehalose levels with SEM are shown of two experiments using three independent transformants of each strain. Statistical analysis was conducted by two-way ANOVA with Bonferroni correction; ns, no significant, *, P < 0.05; **, P < 0.01; ***, P < 0.001****, P < 0.0001.

**Fig 5.**
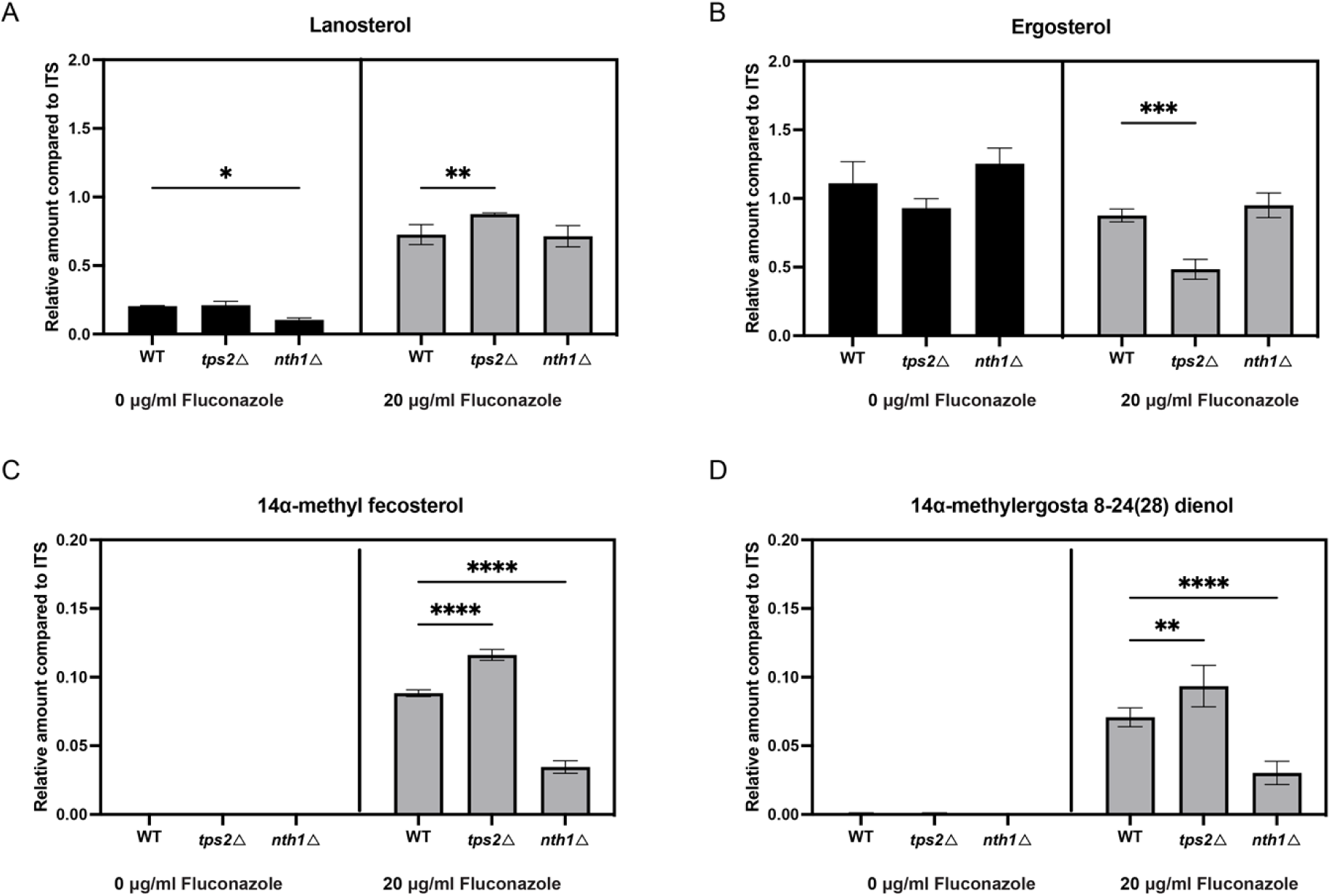
Trehalose metabolism affects the sterol content in *C. glabrata*. Strains were grown in SC glucose medium for 24 h, in the absence or presence of fluconazole. Sterol levels were determined by GC-MS and are displayed for lanosterol, ergosterol, 14α-methyl fecosterol and 14 α -methylergosta-8,24(28)-dienol. The values are calculated relative to the internal standard (ITS; cholestane). Data of the relative amount of sterol compared to ITS, with SEM, are shown from two experiments using three independent transformants of each strain. Statistical analysis was conducted by two-way ANOVA with Bonferroni correction; *, P < 0.05; **, P < 0.01; ***, P < 0.001, ****, P < 0.001.

As Erg11 is the target of fluconazole and the point in the sterol biosynthesis pathway where the flux towards ergosterol or to toxic sterols is defined [39], it seems valid to hypothesize that Erg11 might be the enzyme linking trehalose, or its metabolic enzymes, to ergosterol biosynthesis. We investigated the gene expression levels of *ERG11*, *ERG3*, *ERG6*, *ERG25*, *ERG26* and *ERG27*, involved in the ergosterol biosynthesis pathway in the presence and absence of fluconazole. This will allow us to elucidate whether differences in sterol intermediates are due to different levels of gene expression. Under both conditions, deletion of *NTH1* results in a significant increase in *ERG11* expression, compared to the wild type strain (Fig. 6A). Deletion of *TPS2* results in a lower expression of *ERG11* (Fig. 6A). Furthermore there is a significantly upregulation of *ERG3*, *ERG25* and *ERG26* gene expression under fluconazole-treated conditions (Fig. 6C, D, E) in the *TPS2* deletion strain compared to the wild type strain. Additionally, there was an increase in the expression of *ERG6* and *ERG*27, although these changes were not statistically significant. The lower expression of *ERG11* and the increased expression in the other genes are in accordance with the result that the *tps2Δ* strain exhibits increased levels of toxic sterols and this may contribute to the reduced stress tolerance of this mutant.

**Fig 6.**
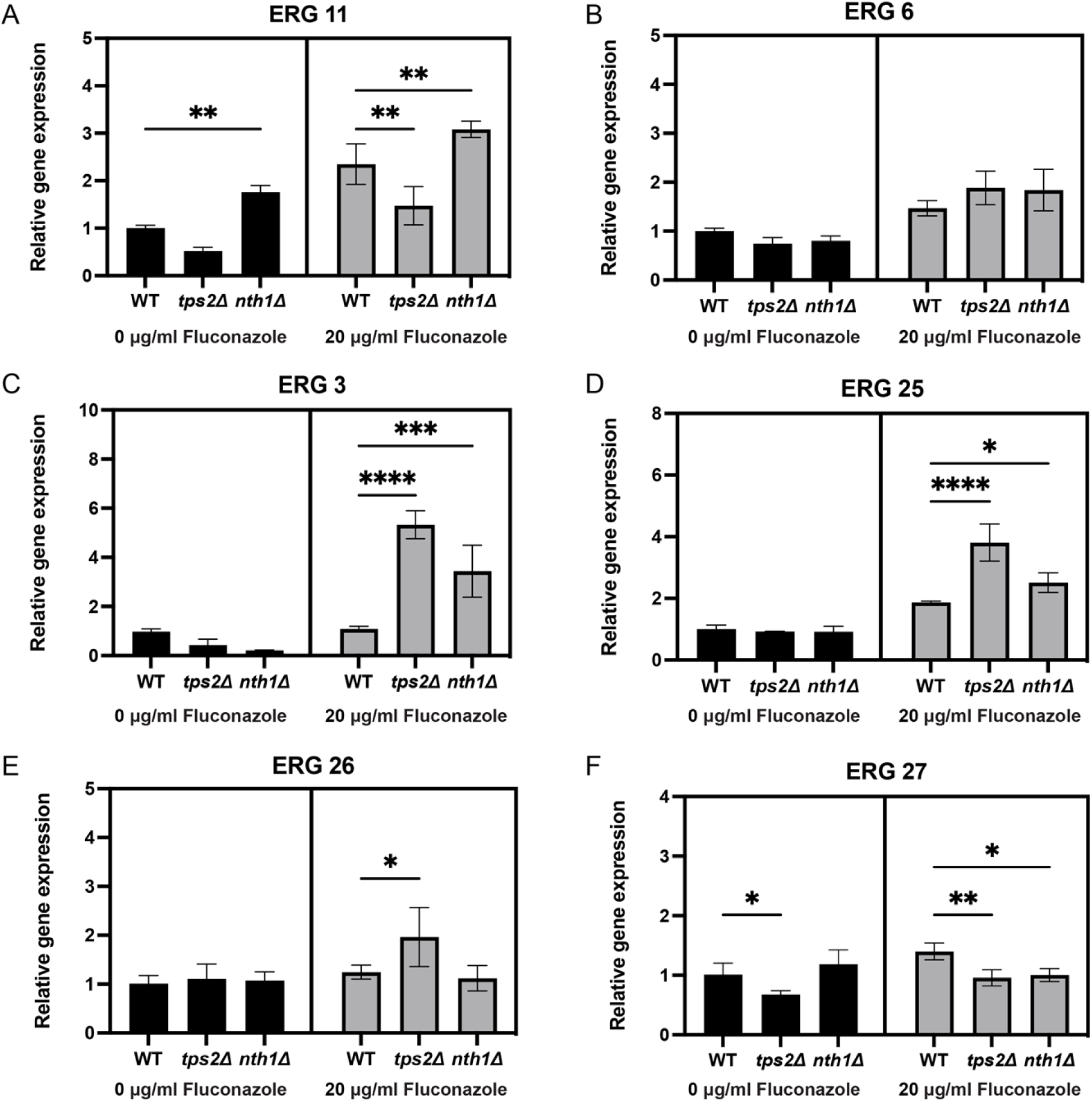
Quantitative RT-PCR determination of the *ERG* gene expression in response to fluconazole stress. Exponentially growing WT, *tps2Δ* and *nth1Δ* strains, in SC glucose medium at 37 °C, were incubated with 20 µg/ml fluconazole for 60 min. Gene expression was analyzed using quantitative reverse transcriptase PCR. Relative gene expression level with SEM are shown from two experiments, each with three independent transformants. The values were calculated relative to the average of the values from the WT untreated samples. Statistical one-way ANOVA analysis of variance with Bonferroni correction was done on the log transformed values (*, P ≤ 0.05, **, P ≤ 0.01).

## DISCUSSION

In this study, mutants lacking the *TPS1*, *TPS2*, and *NTH1* genes in *C. glabrata* were generated and functionally characterized. Our findings underscore the essential role of the *TPS1* gene in glucose utilization as a carbon source, as evidenced by the complete inhibition of growth of the *tps1*Δ strain at 37°C on glucose. Similar to observations in *S. cerevisiae* [40], severe ATP depletion and hexose phosphate accumulation were observed in the *tps1*Δ strain upon glucose addition (data not shown). This effect is attributed to the loss of hexokinase inhibition by T6P, leading to increased glycolytic flux, activation of the protein kinase A pathway and apoptosis [29, 41, 42]. Whereas this phenotype is similar in *S. cerevisiae* and in *C. glabrata*, this is different from what was described in *C. albicans*. Deletion of *TPS1* in *C. albicans* results in a strain that can still grow on glucose at 37 °C but shows growth defects on glucose at higher temperatures [30]. Deletion of the *TPS2* gene results in a strain that accumulates very high levels of T6P under exponential growth conditions. Despite the absence of Tps2, there is still some accumulation of trehalose, a phenotype that was also observed in *C. albicans* and in *S. cerevisiae tps2Δ* mutants, suggesting that some non-specific phosphatases are active under these conditions [43]. A clear difference between *S. cerevisiae* and *C. glabrata* is that under exponential growth conditions on glucose containing medium, *S. cerevisiae* does not accumulate trehalose, whereas we see a clear accumulation in *C. glabrata* (Fig 1C). The absence of trehalose in *S. cerevisiae* is attributed to rapid turnover of trehalose by active trehalase under optimal growth conditions [44]. Based on the very high T6P levels in the *tps2Δ* strain, we assume that the Tps1 enzyme is very active, even under exponential growth conditions in the absence of stress. Deletion of *NTH1* did not result in very high accumulation of trehalose, suggesting that this enzyme is not very active in exponentially growing cells in *C. glabrata*. Deletion of *NTH1* in *S. cerevisiae* increases trehalose levels by 50 to 80 % [45, 46]. A clear observation that distinguishes *C. glabrata* from *S. cerevisiae* is that the human fungal pathogen always accumulates trehalose, even under non-stress conditions. Another distinction is that *C. glabrata* can accumulate very high levels of T6P without showing any growth defect.

In order to have strains that would have high trehalose levels but no accumulation of T6P, we introduced a bifunctional enzyme from *Cytophaga hutchinsonii*, that was previously identified and characterized in the lab [27]. Our hypothesis was that if we expressed the encoding gene into a *tps1Δ* strain, that strain would grow on glucose and would produce high levels of trehalose and no T6P as this should be optimally channeled inside the enzyme. However, we only observe similar trehalose levels as those of the wild type strain but more importantly, there is still very high T6P accumulation (Fig 1D). Therefore, we did not analyze the strains expressing this bifunctional enzyme any further.

Accumulation of trehalose is important to protect cells against various stress conditions. The growth of the *tps2*Δ strain was severely impeded when exposed to heat (42°C), oxidative stress (6 mM H_2_O_2_), and salt stress (1.5M NaCl) (Fig 2). *TPS1* gene expression is increased under these stress conditions (data not shown). Possibly, a further increase in T6P level under these stress conditions may be the underlying reason for the growth defect of the *tps2Δ* strain. Hyper-accumulation of a phosphorylated intermediate, such as T6P, sequesters a large proportion of the cell’s available phosphate. As a consequence, oxidative phosphorylation is limited by the low availability of orthophosphate, leading to low ATP levels and cell death [22, 29, 47]. In contrast, the *NTH1* deletion strain displayed increased tolerance to high temperatures (42°C). However, under salt and oxidative stress conditions, no significant changes in growth conditions were observed compared to the WT strain. This suggests that trehalose accumulation is essential for *C. glabrata* thermotolerance and not for salt or oxidative stress tolerance.

*C. glabrata* is inherently more tolerant to fluconazole and other azoles, compared to other fungal pathogens [48]. The underlying molecular mechanisms are not fully understood [49]. As fluconazole causes stress, we investigated the role of trehalose metabolism in fluconazole tolerance. Our results clearly show that strains with enhanced trehalose accumulation (*nth1*Δ) showed increased resistance and tolerance to fluconazole, whereas strains deficient in trehalose (*tps2*Δ) displayed increased susceptibility to the drug (Fig 3). As T6P levels do not differ greatly between the strains, we conclude that it is the difference in trehalose levels that affect the susceptibility to the drug (Fig 4). As trehalose is known to protect membrane structures under stress conditions, our results may indicate that the indirect effect of fluconazole on membrane structure may be tolerated in *C. glabrata* because of trehalose accumulation.

The primary target of fluconazole is Erg11, an enzyme crucial in the biosynthesis of ergosterol [50] [51]. Our study reveals a notable increase in ergosterol levels associated with the trehalose metabolism pathway, particularly accentuated in the presence of fluconazole (Fig 5). This suggests a protective mechanism whereby trehalose accumulation enables cells to maintain higher ergosterol levels upon fluconazole treatment. Upon detailed examination of sterol profiles in our strains, we observed that the absence of the *NTH1* gene diminishes production of potentially toxic sterols, notably 14α-methylergosta-8,24(28)-dienol [37, 38]. This reduced accumulation of toxic sterols, coupled with enhanced ergosterol production under fluconazole exposure, underscores how higher trehalose levels in the *nth1Δ* strain confer resistance to the drug. *ERG11* plays a pivotal role at the interface of pathways leading to either ergosterol production or toxic sterol accumulation, while *ERG3, ERG6, ERG25*, *ERG26,* and *ERG27* are responsible for both ergosterol and toxic sterol synthesis. Tps2 is essential for normal levels of toxic sterols as our results show an upregulation in expression of genes involved in their production and consequently increased levels of toxic sterols. This suggests that under fluconazole stress, trehalose accumulation protects cells by promoting the flux of the ergosterol pathway and inhibiting the production of toxic sterols. Conversely, the inhibition of Erg11 by fluconazole and the absence of trehalose result in increased levels of toxic sterols. These variations in *ERG11* expression indicate that trehalose influences tolerance by altering *ERG11* expression levels. This modification subsequently affects the metabolic flux towards either ergosterol or toxic sterols. Increased ergosterol production is associated with enhanced tolerance. This hypothesis is strengthened by our results obtained by the *nth1* mutant. Since this mutant has elevated levels of trehalose compared to the WT strain, no increase of toxic sterols is observed. In summary, the trehalose level influences the flux through the toxic sterol biosynthesis pathway by modulating *ERG* gene expression. Specifically, deletion of the *TPS2* gene, which results in lower trehalose levels, is associated with increased toxic sterol levels and decreased ergosterol levels. This emphasizes the role of trehalose in normal functioning of the sterol biosynthesis pathway in the presence of fluconazole. However, the precise mechanism by which trehalose influences ergosterol biosynthesis remains elusive. Whereas in most species it is established that Tps2 is very important (but not completely essential) for trehalose biosynthesis, in *Aspergillus fumigatus* it was shown that trehalose phosphorylases or the regulatory subunit of trehalose biosynthesis, TslA, take over the function to produce trehalose, but no orthologs of these enzymes are found in *C. glabrata* [22, 52]. These authors showed that *Af*Tps2 plays an essential role in cell wall integrity and fungal virulence [22]. Whereas deletion of the *tsla* gene results in lower trehalose levels, the underlying mechanism is not clear as TslA lacks the catalytic site. However, the authors showed that TslA is regulating the activity and subcellular localization of a chitin synthase, providing a link between chitin synthase and trehalose metabolism [52].

Our observations collectively suggest the involvement of trehalose in regulating both the resistance and tolerance to fluconazole, by either a direct effect on the stability of the membrane and/or by altering the sterol profile. Our work again emphasized the potential of trehalose biosynthesis as a promising antifungal drug target [11, 17, 53]. Despite several initiatives, a clear inhibitor of Tps1 or Tps2 is not under clinical investigation, as far as we know. One publication shows an inhibitory effect of N-(phenylthio) phthalimide (NPP) on the Tps2 enzyme of the plant fungal pathogen *Fusarium graminearum* [54] while another study used T6P as a lead compound to develop Tps1 inhibitors [35]. Further research into developing Tps1 or Tps2 inhibitors is necessary to develop a novel class of antifungal drugs, that could work alone or in combination with existing drugs.

## Materials and methods

### Yeast strains, plasmids, primers and media

All strains, plasmids and primers used in this study are listed in <u>Table S1</u> in the supplemental material. Cells were grown at 37°C in SC or YP medium supplemented with 100 mM glucose, unless stated otherwise. YPD medium (1% w/v yeast extract, 2% w/v bacteriological peptone, 2% w/v glucose), YPG medium (3 % v/v glycerol instead of 2 % w/v glucose) and SC medium ( 0.079% w/v complete CSM (MP biomedicals, Santa Ana, CA, USA), 0.17% w/v yeast nitrogen base without amino acids or ammonium sulfate ((NH_4_)_2_SO_4_; Difco), and 0.5% w/v (NH_4_)_2_SO_4_, pH 5.5 (liquid) or pH 6.5 (solid)). SC medium is supplemented with 5 mM glucose, 100 mM glucose or glycerol (3% v/v). RPMI 1640 with L-glutamine (Sigma-Aldrich, Saint Louis, MO, USA) is buffered with 0.165M morpholine propane sulfonic acid (MOPS) to pH 7. For solid media, 2 % w/v agar (Difco) was added.

### Construction of *C. glabrata* Mutant Strains

The trehalose metabolism deletion strains were constructed in the *ATCC2001 his3Δ trp1Δ leu2Δ* background [55]. The wild type strain was transformed with the deletion cassette (the nourseothricin (NAT) marker flanked by FRT sites and a 100 bp region flanking the target gene) by making use of electroporation according to Van Ende et al, 2021 [14]. The deletion cassettes were amplified from the pYC44 plasmid with promoter and terminator sequences of the gene of interest (*TPS1*, *TPS2* and *NTH1*). After transformation, cells were plated on YPG agar medium supplemented with 200 µg/mL nourseothricin. Transformants were checked for insertion of the deletion cassette by PCR and the correct strains were subsequently transformed with the pLS10 plasmid expressing the flippase enzyme to remove the NAT marker (300 µg/ml hygromycin selection). Removal of the NAT cassettes of the transformants was checked via PCR. Finally, the pLS10 plasmid was lost by growth on a nonselective YPG medium and checked by replating on YPG supplemented with 300 µg/mL hygromycin.

The complete coding sequence (CDS) of *ChTPSP* was amplified from the pSal4-ChTPSP plasmid [27]. To express the bifunctional TPS–TPP enzyme, we followed the method outlined in [56], employing expression vectors designed for C-terminal fusions with mCherry fluorescent proteins (pYC56). Subsequently, we substituted the mCherry gene within the pYC56 vector with the *ChTPSP* gene, placing it under the control of a *TEF1* promoter and terminator. To facilitate the expression of the *ChTPSP* gene by replacing the *HIS3* gene, we constructed a replacement cassette by incorporating promoter and terminator sequences derived from the *C. glabrata HIS3* gene into the plasmid capable of expressing the *ChTPSP* gene. Subsequently, we generated a cassette containing the *ChTPSP* gene, a nourseothricin (NAT) marker flanked by FRT sites, flanked by the *HIS3* promoter and terminator sequences, thus forming the novel replacement cassette. Transformation of the wild type, *tps1Δ*, and *tps2Δ* strains through electroporation was performed as previously described.

### Growth Assay

The growth of the mutant strains in both liquid and solid media was monitored over time. For the liquid assay, growth was evaluated by spectrophotometric observation (OD_600_) over time using a Multiskan GO automated plate reader (Thermo Fisher, Waltham, MA, USA). The cultures were diluted in SC medium with glucose (5 mM or 100 mM) and growth was monitored for 36 hours at 37°C. Growth curves were plotted as the average of three biological replicates (= independent mutants). For growth assays on solid medium, a tenfold dilution series of the cultures was spotted on SC plates containing glucose (5 mM or 100 mM). The OD_600_ of the lowest dilution was 0.0001.

### Heat shock, Oxidative and Salt Stress Treatments

The survival of *C. glabrata* cells was determined under various stress conditions using spot assays. Overnight cultures were washed three times with 1x PBS and diluted to OD_600_ of 0.1 in 1x PBS. Next, a tenfold dilution series was made and 5 µL of cells was spotted on SC plates with 2 % w/v glucose. These plates were incubated for 48h at 37°C, 39°C or 42°C for the evaluation of heat stress. Oxidative and salt stress was evaluated on agar containing 6 mM H_2_O_2_ (Sigma-Aldrich) and agar containing 1.5 M NaCl, respectively. Controls were maintained at 37 °C without any treatment.

### Intracellular Trehalose Levels Determination

Exponentially grown cells (25-50 mg) were harvested by centrifugation at 3000 rpm (4 °C) for 5 min. The culture medium supernatant was discarded and the cells were resuspended in 1 ml ice cold Milli-Q water. The cells were broken by vigorously vortexing them in the presence of glass beads (0.75-1 mm) by means of the FastPrep (20s, 6m/s) (MP biomedical™). Subsequently they were immediately transferred to a 95°C water bath for 5 minutes. Afterwards, the sample was centrifuged at 12000 rpm (4 °C) for 5 min and 200 µL of the supernatant was used for analysis by the Shimadzu HPLC system using an Agilent 87 H column at 0.7 mL/min and a RID-20A detector (Shimadzu). High-quality water was used as eluent at a constant flow of 0.6 ml/min at 80°C. Pure trehalose (Sigma) was used as a standard.

### Intracellular Trehalose-6-P Levels Determination

Exponentially grown cells (10-20 mg) harvested by centrifugation at 3000 rpm (4 °C) for 5 min. The cells were extracted as described in Lunn et al [57]. T6P was assayed in extracts by anion-exchange HPLC using a Dionex HPLC system (Sunnyvale, CA, U.S.A.), coupled to an AB Sciex Q-Trap 6500 triple quadrupole mass spectrometer ( as described in [57] with modifications as described in Figueroa et al. 2016 [58].

### RNA extraction and gene expression analysis by qRT-PCR

Cells were grown in SC medium plus 100 mM glucose for 4-5 h until exponential phase at 37°C, while shaking (200 rpm), and fluconazole was added to the cultures to give the final concentrations indicated, and the cells incubated for 60 min. The cells were centrifuged at 14000 rpm (4 °C) for 10 min and washed with 1 ml ice-cold water, resuspended in 1 ml TRIzol (ThermoFisher) and broken using glass beads (0.75-1 mm) in FastPrep machine (20 s, 6m/s). RNA was extracted by respective addition of 360 μl chloroform and 350 μl isopropanol after which three washes with ethanol (70 % v/v) were conducted. RNA was treated with DNase enzyme (New England Biolabs) and converted to cDNA using the iScript cDNA synthesis kit (iScript cDNA synthesis kit; Bio-Rad). Real-time quantitative PCR (qPCR) was conducted using GoTaq polymerase (Promega) and the StepOnePlus real-time PCR device (ThermoFisher). *GAPDH* and *UBC13* were used as reference genes. Data were subsequently analysed using qBasePlus software [59].

### Etest Assays

Fluconazole MICs (Minimum inhibitory concentration) were determined by using Etest strips (BioMérieux) on RPMI (2% w/v glucose) plates. Overnight cultures were washed twice with 1x PBS, adjusted to OD_600_ 0.15 in 1ml 1x PBS, and restreaked on RPMI (2% w/v glucose) plates. Next, a fluconazole-containing gradient strip was placed in the middle of the plate. These plates were incubated at 37°C for 24-72 hours. Subsequently, based on the size of the observed halo, a preliminary assessment of fluconazole tolerance was conducted, and the point of contact between the halo and the strip provided the means to evaluate the MIC_50_ value.

### Broth Microdilution Assays

The broth microdilution assays were based on the Clinical and Laboratory Standards Institute (CLSI) guidelines [60]. Briefly, cells were harvested by centrifugation at 7500 rpm (25 °C) for 1 min from an overnight RPMI (2% w/v glucose) culture, washed three times with 1XPBS, and diluted to around 2500 CFU/ml (verified by plating serial dilution on YPD plates) in RPMI medium. Then round-bottom, UV-sterilized 96-well microtiter plates were used, where all wells were filled with 100 µL diluted cells, 0 to 32 μg/ml fluconazole in 1/2 dilutions, and 80 µL RPMI medium. The plates were incubated at 37°C for 48 hours. The MIC_50_ values were defined as the lowest concentration of the fluconazole that caused ≥50% decrease in optical density at OD_600_. To evaluate the drug tolerance of strains, we generated dose-response curves based on CFU counts for the WT, *tps2Δ*, and *nth1Δ* strains after incubation with fluconazole for 48 hours. The *tps1Δ* mutant was not included as this strain does not grow on glucose. The wells of the broth microdilution assay plate were resuspended, and each culture was diluted and plated on YPD plates. All experiments were conducted with three biological repeats.

### Sterol measurement

Sterols were extracted and analysed based on Morio et al. [61] with some modifications. Cells were grown in SC medium plus 100 mM glucose for 3-4h until exponential phase at 37°C, while shaking (200 rpm), and 20μg/ml fluconazole was added to the cultures and incubated for 120 min. Cells were harvested by centrifugation at 3000 rpm (4 °C) for 5 min, washed twice with MilliQ H_2_O, and a pellet of 20 mg of cells was stored at −80°C. The pellet was resuspended (vortexing for 1 min.) in 300 μl saponification medium (12.5g KOH in 18ml MilliQ H_2_O diluted to 50ml with 98% ethanol), transferred to a capped glass vial and incubated for 1h in a shaking water bath at 80°C. Sterols were extracted by adding 100 μl MilliQ H_2_O and 400 μl hexane, including 1 μl of 5 mg/ml 5-a-cholestane as internal standard (Sigma, 47124), followed by vortexing for 3 min, 20 min. phase separation and collection of 350 μl of the top (hexane) layer. A second extraction fraction was collected by adding 600 μl hexane, vortexing for 3 min., 20 min. phase separation and collection of 550 μl of the top (hexane) layer. The two collected hexane fractions were combined and dried using vacuum centrifugation (Automatic Environmental SpeedVac System AES2010) for 30 min at room temperature. Sterol extracts were re-dissolved in 60 μl hexane and derivatized by adding 10 μl of silylating mixture (Sigma, 85432), short vortexing, and incubation at room temperature for 1 h. Derivatized extracts were shortly centrifuged to precipitate potential debris and 50 μl of the extract was transferred to a smaller insert glass tube for GCMS analysis. The samples were analysed using a Thermo Scientific gas chromatography-mass spectrometer (Trace 1300 - ISQ QD equipped with a TriPlus RSH autosampler and a Restek Rxi-5ms capillary GC column (30 m x 0.25mmID)). Helium was used as carrier gas with a flow rate of 1.4 ml/min. Injection was carried out at 250°C in split mode after 1 min and with a ratio of 1:10. The temperature was first held at 50°C for 1 min and then allowed to rise to 260°C at a rate of 50°C/min, followed by a second ramp of 2°C/min until 325°C was reached; that temperature was maintained for 3 min. The mass detector was operated in scan mode (50 to 600 atomic mass units), using electron impact ionization (70 eV). The temperatures of the MS transfer line and detector were 325°C and 250°C, respectively. Sterols were identified by their retention time relative to the internal standard (cholestane) and specific mass spectrometric patterns using Chromeleon software (version 7). Abundance was calculated relative to the internal standard, comparing the relative peak areas of the compounds. Sterol extraction and analysis of each strain was performed in duplicate (technical repeats) on individually cultured strains.

## Acknowledgments

This work was supported by a grant from the Fund of Scientific Research Flanders (FWO # G0C0622N) and by the Max Planck Society (R.F and J.E.L.). QZ acknowledges the Overseas Study Program of Guangzhou Elite Project (nr: S.J. [2020] No. 2) for a PhD fellowship.

## Declaration of interests

The authors have no interests to declare.

## Notes

### Competing Interest Statement

The authors have declared no competing interest.

